# Analysis of Gene Expression Changes upon Topobexin Treatment and TOP2B-knockout in hiPSC derived cardiomyocytes

**DOI:** 10.64898/2026.03.09.710520

**Authors:** Veronika Kerestes, Ian Cowell, Anna Jirkovska, Mushtaq Khazeem, Galina Karabanovich, Iuliia Melnikova, John Casement, Jan Kubes, Tomas Simunek, Jaroslav Roh, Matthew Schellenberg, Adam Creigh, Chunbo Yang, Majlinda Lako, Lyle Armstrong, Caroline Austin

**Affiliations:** Newcastle University, Faculty of Medical Sciences, UK; Charles University, Faculty of Pharmacy, Hradec Kralove, Czech Republic; Mayo Clinic, Rochester, USA; Mustansiriyah University, Iraq

**Keywords:** TOP2, TOP2B, gene expression, cardiomyocyte, hiPSC

## Abstract

The role of DNA topoisomerase II beta (TOP2B) in cardiomyocyte differentiation is poorly understood. To address this, Human induced pluripotent stem cells (hiPSC) were differentiated into cardiomyocytes (CM) that are wildtype or contain a genomic deletion of Topoisomerase 2B (BKO). Both WT and BKO hiPSC could be induced to differentiate into sheets of beating cardiomyocytes. BKO hiPSC take slightly longer to differentiate into sheets of beating CM than WT iPSC. RNA was prepared from both undifferentiated and differentiated WT and BKO hiPSC. RNA seq was used to examine gene expression changes when the WT and BKO hiPSC were differentiated into CM. Gene expression changes following differentiation of BKO cells were largely similar to those in WT cells. In addition, the differentiated WT CM were treated with dexrazoxane (ICRF-187), a TOP2 catalytic inhibitor that targets both TOP2A and TOP2B, or topobexin, a new TOP2B selective catalytic inhibitor. Topobexin inhibition partially phenocopied a TOP2B deletion and thus providing an alternative to TOP2B gene knockout in many cell lines. In future, hiPSC derived CM with and without TOP2B and inhibition by topobexin ex vivo CM could be used to study anthracycline-induced cardiotoxicity and to screen for cardioprotectants.

**Highlights:**

- Used CRISPR-Cas9 to delete TOP2B from hiPSC
- Produced beating cardiomyocytes from both WT and TOP2B null hiPSC
- Transcriptome analysis of WT and TOP2B null hiPSC and derived cardiomyocytes
- RNA seq showed he specific TOP2B inhibitor topobexin largely phenocopies TOP2B gene inactivation in iPSC derived cardiomyocytes.
- Topobexin inhibition could be used as an alternative to a TOP2B gene knockout in many different cell types, speeding up the analysis of the function of TOP2B.

## Introduction

DNA topoisomerase II beta (TOP2B) is an essential nuclear enzyme that modulates DNA topology by introducing transient double-strand breaks. Unlike the alpha isoform (TOP2A), which is primarily active in proliferating cells, TOP2B is expressed ubiquitously, including in post-mitotic cells, such as neurons and cardiomyocytes. TOP2B facilitates the resolution of topological constraints during transcription and participates in long-range chromatin remodelling and gene activation [1,2]. TOP2B is involved in regulating transcription [3] but is not essential for cell viability [4–6]. However, TOP2B is required in some differentiation pathways including neuronal and B cell differentiation [7–10].

Using TOP2B^-/-^ embryonic fibroblast cell lines [11] showed a connection between TOP2B and the prevention of anthracycline cardiotoxicity with dexrazoxane (ICRF-187). A link between TOP2B and anthracycline induced cardiotoxicity was confirmed by [12] using an *in vivo* conditional murine cardiomyocyte-specific knockout study that firmly established TOP2B as the causal driver of anthracycline-induced cardiotoxicity [13]. Two studies have used iPSC derived cardiomyocytes to investigate the role of TOP2B on anthracycline-induced cardiotoxicity: one study used CRISPR-Cas9 deletion of TOP2B [14] and a second used siRNA knockdown of TOP2B [15]. These studies showed that deletion or knockdown of TOP2B reduced the loss of viability of the cells following exposure to doxorubicin. Furthermore, Saroj *et al*, showed that siRNA knockdown was more effective than (ICRF-187) treatment, and they suggested siRNA mediated TOP2B inhibition could provide a way to mitigate doxorubicin cardiotoxicity. However, methods for effective delivery of siRNA to cardiomyocytes *in vivo* have not yet been established, and a small molecule drug would be more clinically achievable approach. Notably, topobexin a specific TOP2B inhibitor has recently been reported [16].

In this study, we developed hiPSC with CRISPR-Cas9 deletion of TOP2B (BKO) and differentiated them into beating cardiomyocytes. In the selected clone (line 18D) we performed transcriptomic profiling to evaluate gene expression changes before and after differentiation. In addition, we compared the transcriptional effects of TOP2B genetic deletion with the effects of its pharmacological inhibition by the non-isoform-selective inhibitor ICRF-187 and by a novel TOP2B selective catalytic inhibitor topobexin [16]. By comparing genetic inactivation with pharmacological inhibition, we aimed to define the role(s) of TOP2B in cardiomyocyte differentiation and transcriptional regulation for exploring how modulation of TOP2B activity may influence cardiotoxic responses and broader aspects of cardiac biology.

## Materials and methods

### Cell culture

Human iPSCs were derived from unaffected fibroblasts purchased from Lonza a commercial supplier. Undifferentiated hiPSC cells were maintained in Geltrex (Thermo-Fisher)-coated wells of 6-well plates in mTeSR1 medium (StemCell Technologies). For the first 24 hours after plating the medium contained 10 µg/ml ROCK inhibitor (Y27632, StemCell Technologies), after which the medium was replaced with mTeSR1 without ROCKi.

### CRISPR-Cas9 deletion of TOP2B

TOP2B was deleted from hiPSC using CRISPR-Cas9. The gRNA targeting exon 1 (See Supplementary Fig. 1A) was cloned into plasmids as described in [9]. For transfection, 2 × 10⁶ cells were centrifuged (115 × g, 3 min), resuspended in electroporation solution (Amaxa Human Stem Cell Nucleofector Kit, Lonza), and transferred to a cuvette. Plasmids were added, and nucleofection was performed using the Amaxa Nucleofector™ II, program A-23. Cells were then plated on a Geltrex-coated 6-well plate, and conditioned mTeSR1 (Stemcell Technologies) medium (1:1 fresh and pre-used, filtered through a 0.22 µm PES filter) was added. After 48 hours, cells were detached with Accutase (Biosera) and mixed with 2 ml FBS. Clumps were removed using a 40 µm sieve, and single-cell suspensions were transferred to flow cytometry tubes. Hoechst 33342 (Thermo Fisher Scientific, 0.1 µl/ml) was added for nuclear staining. Cells were sorted based on GFP expression into Geltrex-coated 96-well plates containing 100 µl conditioned mTeSR1 medium with ROCK inhibitor (10 µg/ml). After 24 hours, the medium was replaced with fresh mTeSR1 and changed every other day. Visible colonies appeared after about one week, and by four weeks, genotyping could be performed. Screening for TOP2B knockout clones was performed using PCR genotyping and confirmed by DNA sequencing and Western blotting (see supplementary Figures 1B & 1C).

### Immunodetection of TOP2B

Pellets from 1×10⁶ hiPSC were resuspended in extraction buffer (10 mM MgCl₂, 50 mM Tris-HCl pH 7.4, 4 mM DTT) with 0.05% protease inhibitors (Proteoloc Expedeon, Abcam), then treated with 0.25% SDS and 50 µg/ml DNase I (Thermo Fisher Scientific). After incubation on ice (10–30 min), an equal volume of solubilization buffer (2% SDS, 20% glycerol, 5% β-mercaptoethanol, 0.6 M Tris-HCl pH 7.2) was added, and samples were heated at 68 °C for 10 min. Each sample was mixed with 20 µl of sample buffer, heated at 68 °C for 5–10 min, and loaded onto 10-well gradient SDS-PAGE gels. Electrophoresis was run in Tris-HEPES-SDS buffer (NuSep) at 120 V for 80 min. Proteins were transferred to nitrocellulose membranes using wet transfer (25 mM Tris, 192 mM glycine, 20% methanol) on ice at 110 V and 500 mA for 70 min. Membranes were blocked overnight at 4 °C in 5% milk in 0.1% TBS-T. The next day, they were incubated with primary anti-TOP2B antibody (MAB6348, 1:500, R&D Systems) and anti-β-actin antibody (NB600-501, 1:5000, Novusbio) for 1 h at room temperature, then washed with 0.1% TBS-T (2× 1 min, 1× 10 min, repeated twice). After incubation with HRP-conjugated secondary antibody (NXA931, 1:5000, GE Healthcare) for 1 h at room temperature, membranes were washed again using the same protocol. Signal was detected using Clarity ECL substrate (Bio-Rad) and imaged with a C-DiGit blot scanner (Li-Cor) (supplemental figure 1C).

### Cardiac Differentiation of hiPSC

Differentiation was initiated at 70% confluence derived from [17] by replacing mTeSR1 (Stemcell Technologies) medium with RPMI 1640 (Gibco) supplemented with B27 without insulin (Thermo Fisher Scientific), 10 µM CHIR 99021 (Tocris Bioscience) and 10 ng/ml Activin A (Tocris Bioscience). Cells were then treated following the protocol outlined in Table 1. Spontaneous contractility appeared after 8 days or later.

**Table 1.** hiPSC differentiation protocol.

| Day | Medium | Growth Factors |
| --- | --- | --- |
| 1 | RPMI 1640 + B27 without insulin | 10 μM CHIR 99021, 10 ng/ml Activin A |
| 2 | RPMI 1640 + B27 without insulin | None |
| 4 | RPMI 1640 + B27 without insulin | 5 μM IWP2 |
| 6 | RPMI 1640 + B27 without insulin | None |
| 8 | RPMI 1640 + B27 with insulin | None |
| Every other day | RPMI 1640 + B27 with insulin | None |

### Sample preparation for differential gene expression analysis

Cells were cultured as in the previous CM differentiations, using 6 well plates with independent differentiations in each well. Total RNA was prepared using an RNase easy mini kit (Qiagen). Samples were prepared from undifferentiated or differentiated cells (beating sheets of CM cells). Additionally, total RNA samples were prepared form differentiated CM cells after treatment with ICRF-187 (100 µM for 3 hours immediately prior to cell lysis) or treatment with topobexin (10 µM for 3 hours immediately prior to cell lysis). Four replicas were prepared for each condition. For each replicate RNase easy prep was prepared from a single well of a 6-well plate. RNA libraries for RNA-seq were prepared using Illumina Stranded mRNA Prep, Ligation library preparation protocol. Single-end sequencing was performed using an Illumina NovaSeq 6000 sequencer. Transcript-level quantification was performed using Salmon [18]. Subsequent analysis steps employed R software (version 4.3). PCA analysis employed DESeq2 [19]. The GEO accession number for RNA-seq data reported here is GSE262148.

## Results

### Generation of knockout hiPSC

TOP2B null hiPSC clones were generated from wild-type hiPSC cells previously reported (AD1/WT1) [20] using CRISPR-Cas9 as previously described for SH-SY5Y neuroblastoma cells using TOP2B exon 1 guide RNA 6 [9] (Fig. S1A). TOP2B knockout was confirmed by DNA sequencing and by western blotting the selected clones. Data for clone 18D is shown in supplemental Figure S1B and S1C). TOP2B null lines (BKO) appeared phenotypically indistinguishable to the WT with identical appearance and growth properties. However, differential gene expression comparisons between the undifferentiated TOP2B null cell line and the WT cell line did reveal some transcriptional changes. These are highlighted in a volcano plot in supplementary Fig. S1D. With a cut-off of a twofold change in transcript abundance, 101 genes were significantly down regulated genes and 426 up regulated in the TOP2B null hiPSC line.

### Differentiation of WT and knockout iPSC

hiPSC were differentiated using a protocol derived from that reported by [17]. Both WT and BKO hiPSC differentiated efficiently and reproducibly using this protocol. Four biological replicates for the WT hiPSCs and five biological replicates for the BKO hiPSCs. However, differentiation to the endpoint of beating sheets of CM took approximately 1 day longer for the TOP2B null cells after the replacement of mTeSR1 with RPMI 1640 + B27 + GSK3 inhibitor (CHIR 99021) + Activin A (see Fig 1A). The time until beating for each cell line and each replicate is shown in Figure 1A. The mean time to beating for WT CM was 9 days and for BKO CM it was 10.5 days.

**Figure 1.**
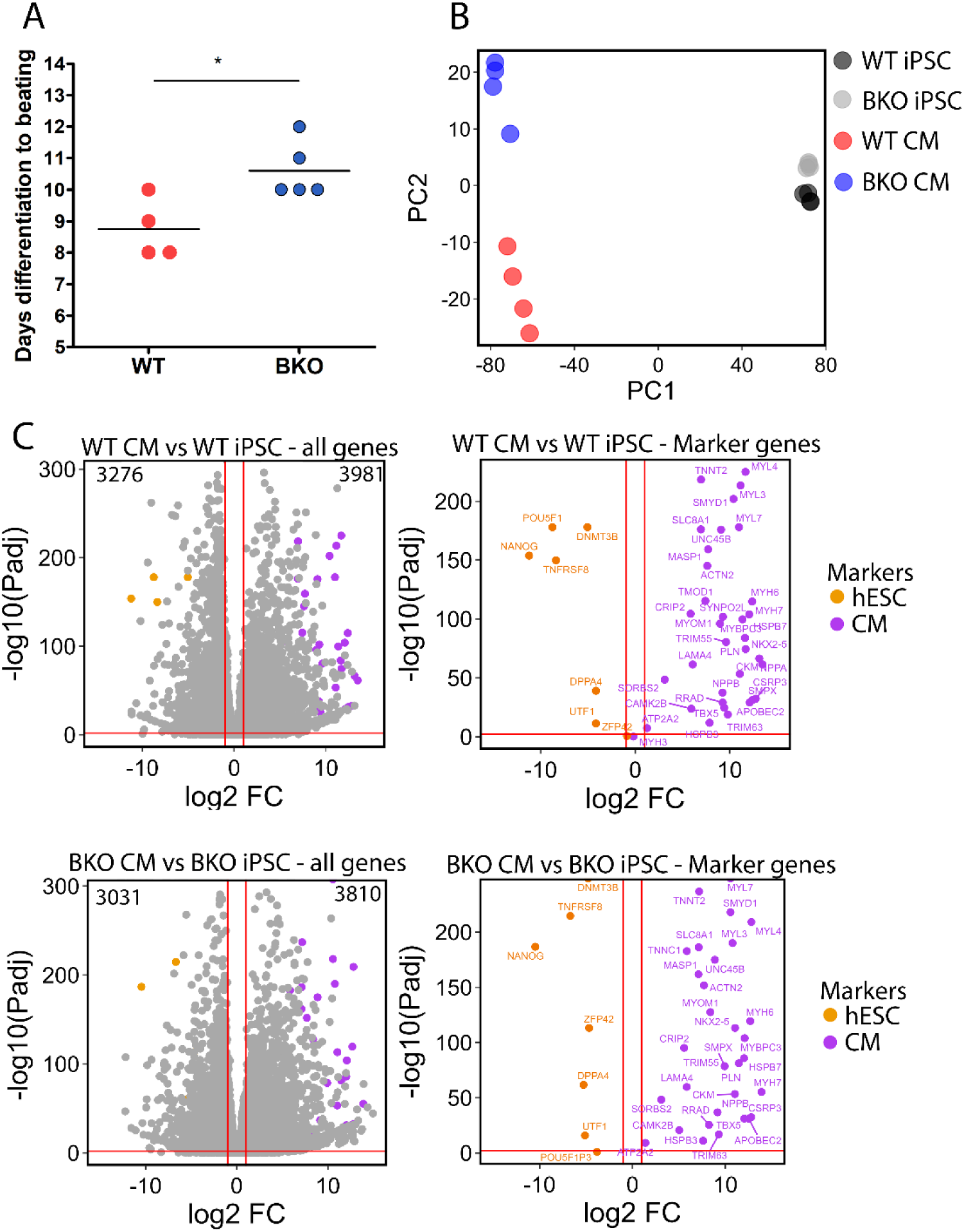
Wild type and TOP2B null hiPSCs can efficiently differentiate into beating cardiomyocytes. **(A)** The differentiation protocol resulted in the conversion of monolayers of wild type (WT) or TOP2B null (line 18D, BKO) hiPSC cells into layers of beating cardiomyocytes in 8-12 days. However, on average this took 1 day longer for BKO than WT hiPSC cells. Each spot represents a separate hiPSC to CM differentiation experiment, four biological replicates for WT and five biological replicates for the BKO. Line = mean. Statistical significance was determined using a two-tailed unpaired t-test, * - *P* < 0.05. **(B)** Primary component analysis of RNA-seq analysis comparing WT and TOP2B null (BKO) hiPSC and derived CM (quadruplicate biological replicas). **(C)** Volcano plots to visualise gene expression changes upon differentiation of WT and BKO hiPSC cells to CM. Red lines represent cutoffs of > 2-fold change in transcript abundance with a *P*adj of <0.05. Numbers in the corners of the plots highlight the number of up or down regulated protein-coding genes in the CM compared to the hiPSC cells, using the above cutoffs.

### Differential expression analysis confirmed the differentiation of WT and BKO cell lines-increased expression of cardiomyocyte marker genes

A comparative exome analysis was performed in WT hiPSC and hiPSC null for TOP2B (BKO hiPSC) that were generated by targeted gene knockout. RNA-sequencing was performed on four biological replicates for each condition (undifferentiated WT hiPSC, undifferentiated BKO hiPSC, WT differentiated into cardiomyocytes (WT CM), BKO differentiated into cardiomyocytes (BKO CM). Figure 1B shows principal component analysis for the four conditions. This shows the biggest difference is between the undifferentiated cell lines and the differentiated cell lines. The undifferentiated cell lines clustered together away from the differentiated cells. As expected, the largest difference in gene expression profiles was between the hiPSC and derived CM cells of both genotypes (PC1); but notably the four WT CM replicas cluster separately from the four BKO CM samples (PC2). This is consistent with a largely shared pattern of transcriptional changes upon differentiation to the CM state, but with some differences between the WT and BKO cells in this respect. The differential gene expression between the undifferentiated and differentiated cell lines is illustrated in the volcano plots in Figure 1C. Comparison of the differentiated and undifferentiated cell lines showed that similar numbers of genes are up or down regulated upon differentiation in WT or BKO cell lines (red lines indicate 2-fold change in gene expression). Pluripotency (human embryonic stem cell, hESC) and cardiomyocyte marker genes have previously been identified in global expression profiles of highly enriched cardiomyocytes derived from human embryonic stem cells in [21] (Table 2). As expected, cardiac marker genes were upregulated in CM cells of both genotypes. These genes included *ACTN2* (alpha actinin 2), *MYL7* (Myosin light chain 7) and *TNNT2* (troponin 2, cardiac type). Similarly, markers of pluripotency including *NANOG*, *DPPA4* and *DNMT3B* were downregulated in both WT and BKO CM cells Fig. 1C.

**Table 2.** Cardiomyocyte (CM) and pluripotency (hESC) marker genes, from [21].

| Gene symbol | Type |
| --- | --- |
| ACTN2 | CM |
| APOBEC2 | CM |
| ATP2A2 | CM |
| CAMK2B | CM |
| CKM | CM |
| CRIP2 | CM |
| CSRP3 | CM |
| HSPB3 | CM |
| HSPB7 | CM |
| LAMA4 | CM |
| MASP1 | CM |
| MB | CM |
| MYBPC3 | CM |
| MYH6 | CM |
| MYL3 | CM |
| MYL4 | CM |
| MYL7 | CM |
| MYOM1 | CM |
| PLN | CM |
| PPP1R12B | CM |
| PPP1R14C | CM |
| RRAD | CM |
| SLC8A1 | CM |
| SMPX | CM |
| SMYD1 | CM |
| SORBS2 | CM |
| SRD5A2L2 | CM |
| SYNPO2L | CM |
| TMOD1 | CM |
| TNNC1 | CM |
| TNNT2 | CM |
| TRIM55 | CM |
| TRIM63 | CM |
| UNC45B | CM |
| NPPA | CM |
| NPPB | CM |
| TBX5 | CM |
| MYH7 | CM |
| NKX2-5 | CM |
| TNNC1 | CM |
| NANOG | hESC |
| ZFP42 | hESC |
| POU5F1 | hESC |
| TNFRSF8 | hESC |
| DPPA4 | hESC |
| UTF1 | hESC |
| DNMT3B | hESC |
| LIN28 | hESC |

### Differences in gene expression between WT and BKO CM

Although there were a similar number of gene expression changes when either WT or BKO hiPSC differentiated into beating CM (Fig. 1C), differential gene expression comparisons between the differentiated BKO CM and WT CM showed 530 genes were down regulated and 760 up regulated in the BKO CM compared to WT CM (Fig. 2A), so there are slightly more up-regulated than down-regulated genes. None of the CM markers were affected significantly comparing WT to BKO CM, nor were the hESC genes, except for *ZFP42*, which was more abundant in the BKO CM compared to WT. However, this gene is expressed at a much higher level in the BKO hiPSC than in their WT equivalents (see Supplementary Table 1). Comparing genes whose expression changed during differentiation (>= 2-fold change, *P*adj <0.05), the majority of changes (5531) were shared between both genotypes. However, 1672 and 1267 genes were differentially expressed only in WT or BKO respectively during differentiation to CM (See supplementary Figure S2).

**Figure 2.**
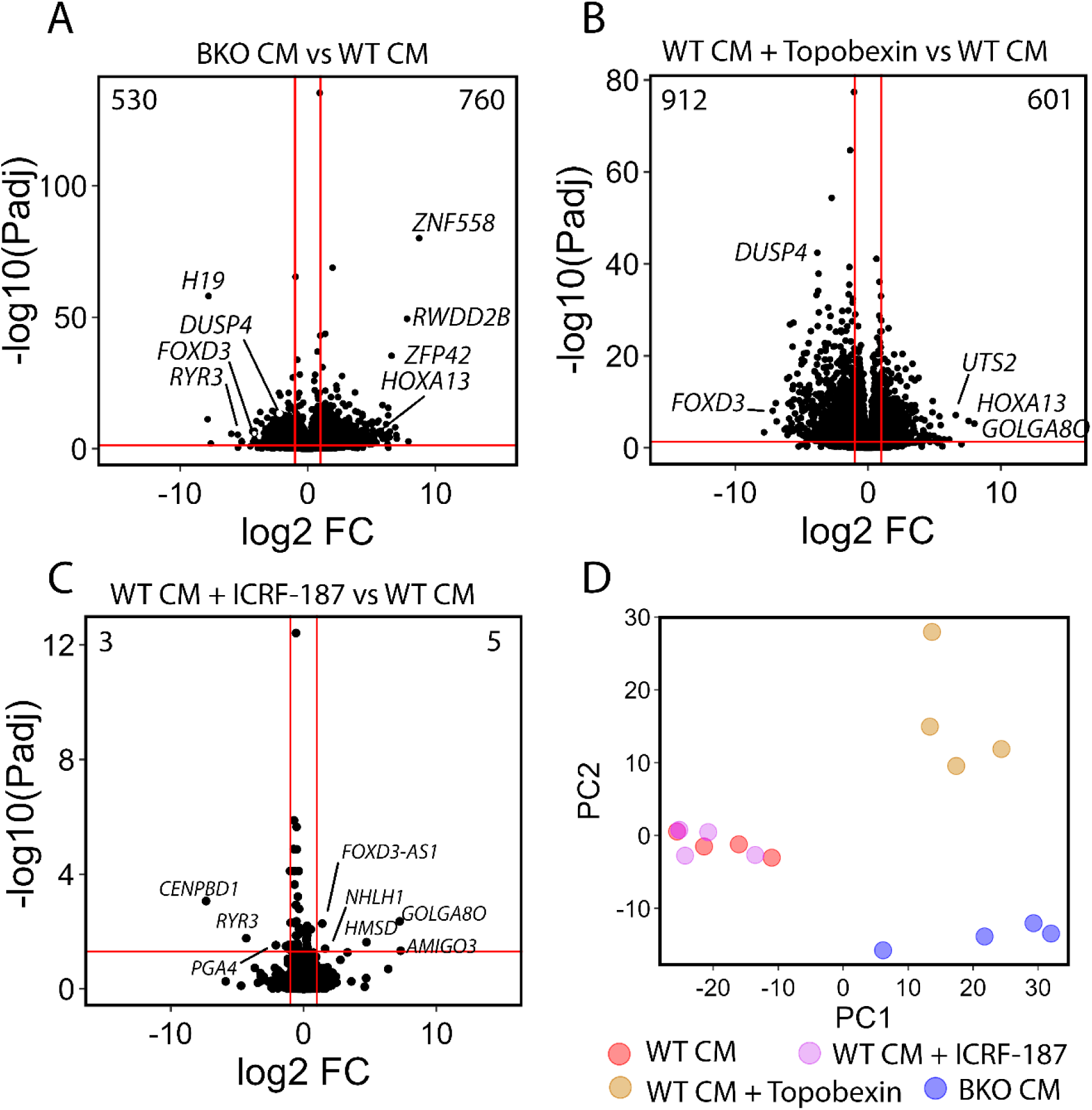
Transcriptome comparison of WT and BKO hiPSC-derived CM cell, and effect of TOP2 inhibitors topobexin and ICRF-187. **(A)** Volcano plot illustrating the number of genes up- or down regulated in BKO versus WT CM. **(B)** Effect of topobexin gene expression (transcript abundance) in WT CM. **(C)** Effect of ICRF-187 on gene expression (transcript abundance) in WT CM. For volcano plots, cut-offs (red-lines) were as in Fig. 1. Representative genes are highlighted. **(D)** Primary component analysis of transcript abundance comparing in four conditions, WT and TOP2B null (BKO) CM, WT CM treated with topobexin and CM treated with ICRF-187 (quadruplicate biological replicas).

### Effect of topoisomerase II inhibition on gene expression in the WT CM

To compare topoisomerase II inhibition with TOP2B knockout WT CM were treated with two catalytic inhibitors: topobexin [16], a recently developed small molecule with selectivity for TOP2B (10 µM, 3hr, Fig 2B) or dexrazoxane (ICRF-187), a topoisomerase II inhibitor used clinically as a cardioprotectant (100µM, 3hr, Fig 2C) using concentrations and exposure times derived previously. Figure 2B & C show volcano plots of the differential gene expression patterns following treatment with topobexin or ICRF-187. Topobexin had a large effect, with 912 down regulated and 601 upregulated genes. In contrast, ICRF-187 had very little effect on gene expression in WT CM. None of the CM or hESC marker genes were significantly affected, and even taking the whole transcriptome, only eight genes were significantly altered in ICRF-187 treated cells (see Fig 2C). A table of relative gene expression values (TPM values) for each condition is given in (Supplementary Table S1). Notably, all of the genes whose expression was significantly altered by ICRF-187 were expressed at relatively low levels (Supplementary Table 1). Transcripts corresponding to one of the genes upregulated by ICRF-187 (*GOLGA8O)* were also increased in abundance in BKO CM and topobexin-treated WT CM (Fig. 2, Supplementary Table 1). Genes with a large increase in transcript abundance in the BKO CM included *ZNF588, RWDD2B, HOXA13* and the previously mentioned *ZFP42* and *GOLGA8O.* Of these selected genes, *HOXA13* and *GOLGA8O* were also elevated in topobexin-treated cells, while *ZNF588, RWDD2B* and *ZFP42* were not. Conversely, transcript abundance for genes such as *UTS2* was increased only in the topobexin-treated CM. Looking at selected down-regulated transcripts, one of the most under expressed transcripts in the BKO cells, *H19*, is not changed in abundance in topobexin treated CM, while *DUSP4* and *FOXD3* were downregulated in both BKO and topobexin-treated cells.

A principal component analysis was carried out on the four differentiated conditions (untreated WT CM, BKO CM and WT CM cells treated with ICRF-187 or topobexin). For ICRF-187 treated WT CM, all four ICRF-187-treated CM replicates clustered together with the untreated WT CM, consistent with the limited transcriptional changes highlighted in Fig. 2C. In contrast, topobexin treated WT CM and BKO CM were separated from the untreated/ICRF-187 cluster by PC1, although they were separated from each other along the PC2 axis (Fig. 2D). About half of the genes whose expression was changed in the TOP2B null CM compared to WT, were also changed in the topobexin treated WT CM – consistent with the PCA analysis (Fig. S2 B). However, fewer than half of the topobexin affected genes were also altered in the knockout (Fig. S2B). Of the genes whose expression changed in both conditions (WT vs BKO CM cells and WT CM cells vs WT CM cells + topobexin) we observed a significant correlation in the size and direction of the change in transcript abundance (Pearson corelation coefficient = 0.88, Fig. 3). This suggests that topobexin partially phenocopies the BKO cells.

**Figure 3.**
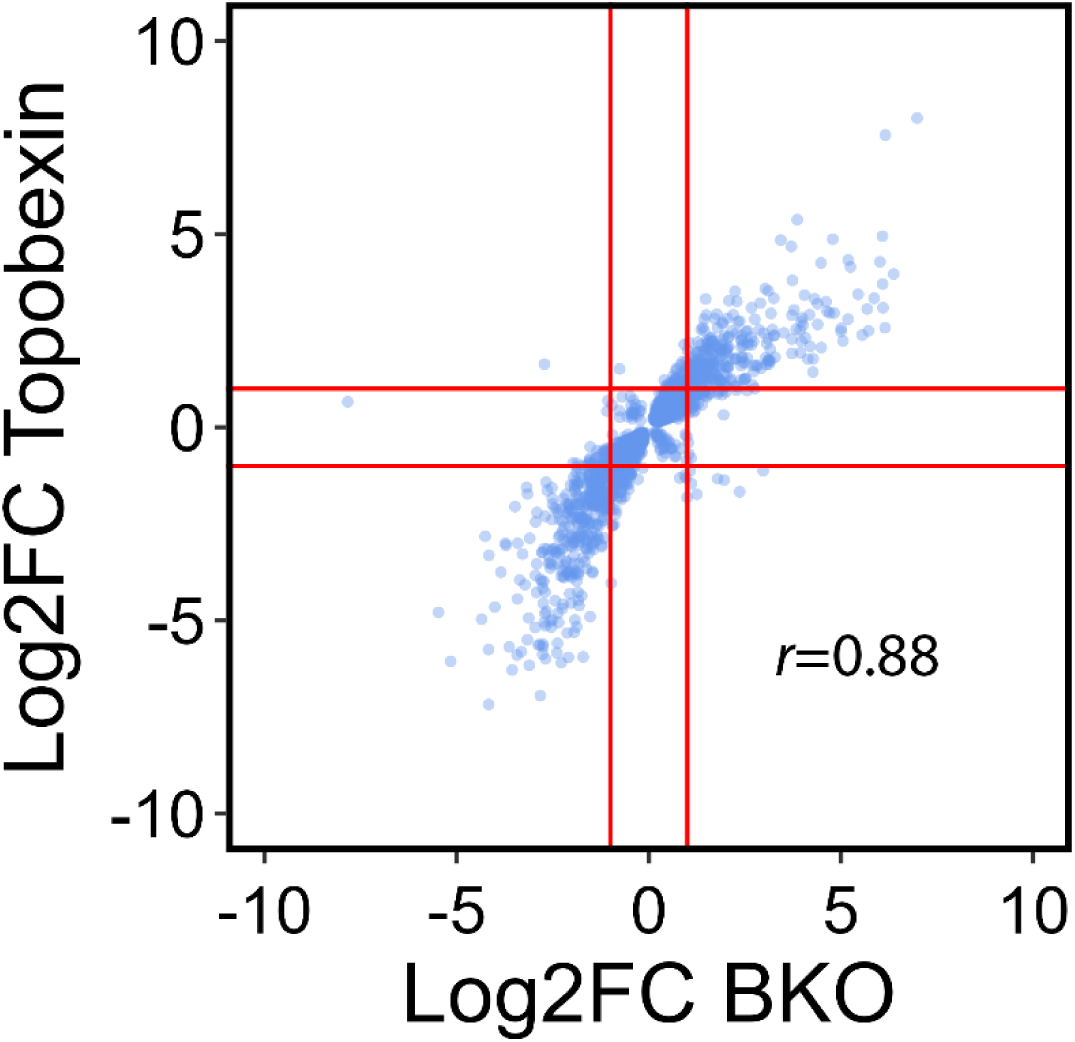
Comparison of gene expression changes in WT vs BKO CM and topobexin treated vs untreated CM. Differential gene expression tables were filtered to include only genes that change significantly (*P*adj <0.05) in expression in both TOP2B null CM vs WT cells and after treatment of WT CM with topobexin. The gene list was further filtered to only include protein-coding genes. Log2 FC values are plotted on the x and y axes. *r* = Pearson corelation coefficient.

### Gene set enrichment analysis

We carried out Gene Set Enrichment Analysis using g:GOSt [22] in order to determine whether any specific biological pathways or process where disproportionately represented in the sets of genes whose transcript abundance significantly changed in either BKO CM or topobexin-treated CM compared to untreated WT CM. A large number of Biological Processes (BP) terms were highly significantly enriched for both gene sets. The BP terms most enriched related to cellular and tissue development and differentiation. The top 40 scoring BP terms are plotted by significance value in Supplementary Fig. 3, and the entire data set is present in Supplementary Table 2.

### Effect of removal of TOP2B on gene expression of long genes in the CM

TOP2B has been reported to be involved in the transcription of long genes [3]. For example in SH-SY5Y cells, the longest genes were more likely to be downregulated in TOP2B null compared to the WT cells [9]. To determine if this also occurs in cardiomyocytes an equivalent analysis was carried out for the WT versus BKO CM and for control versus topobexin-treated cells. As we observed previously, down-regulated transcripts were slightly more common in the upper quartile of gene lengths (Table 3), although this effect was not as pronounced as observed when comparing WT to TOP2B null SH-SY5Y cells. In WT vs BKO CM, downregulated genes were least abundant in the middle two quartiles, and this pattern was also present in topobexin-treated cells. Taken together the data in Fig. 2, Fig. 3 and Table 3 support the idea that topobexin partially phenocopies TOP2B gene inactivation in hiPSC-derived cardiomyocytes.

**Table 3.** Relationship between gene length and probability of a gene being down regulated in TOP2B null or topobexin-treated CM (Cut-offs, *P*adj <0.05, Log2FC < −1)

| Gene length quartile | WT v BKO (TOP2B null) |  | WT vs WT + Topobexin |  |
| --- | --- | --- | --- | --- |
|  | Number of genes | Percentage of downregulated genes | Number of genes | Percentage of downregulated genes |
| Q1 | 152 | 28.7 | 239 | 26.5 |
| Q2 | 108 | 20.4 | 198 | 22.0 |
| Q3 | 96 | 18.1 | 184 | 20.4 |
| Q4 | 174 | 32.8 | 280 | 31.1 |
| Total | 530 | 100 | 901 | 100 |

## Discussion

Classical *in vitro* cardiac models often rely on cell lines or primary cardiomyocytes derived from non-human species such as rodents. *In vitro* models of cardiomyocytes are indispensable tools for studying cardiac physiology, molecular signalling, and responses to external stimuli. These models support both basic research and applied biomedical studies, including drug development and safety testing. However, while these systems are accessible and well-established, they may not fully replicate human-specific molecular mechanisms, electrophysiological properties, or gene regulation patterns. Cross-species differences can limit the translational value of findings, particularly when investigating subtle regulatory processes or drug responses that are specific to human biology. Immortalized cardiomyocyte-like cell lines and rodent primary cardiomyocytes present additional limitations. For example, cell lines frequently exhibit altered signalling pathways and an immature cardiac phenotype. Primary cardiomyocytes, although more physiologically relevant, are difficult to isolate in large numbers, are short-lived in culture, and generally cannot be propagated. These challenges underscore the need for more physiologically relevant and genetically tractable human cell-based models. Many diseases or pathophysiological mechanisms seen in human medicine are difficult to model in experimental cellular or animal systems reproducibly and directly translatable into clinical situation [23].

Human derived hiPSCS have numerous advantages over animal models or immortalized cell lines. They can be differentiated into a variety of cells including cardiomyocytes [24]. Also, they can be used in high-throughput screening [25], patient-specific research or personalised medicine [26], and the response to drugs can be explored in cardiac organoids [27]. Thus, hiPSC offer a compelling alternative. Derived from adult somatic cells, hiPSC-derived cardiomyocytes (hiPSC-CM) provide a renewable, human-specific model system that allows for genetic manipulation and disease modelling in a controlled setting. They can reproduce many key structural and functional features and may be used to investigate developmental processes, gene regulation, and cellular responses in a human context.

TOP2B has also been implicated in the mechanism of anthracycline-induced cardiotoxicity, although generally the role of TOP2B in human cardiomyocytes remains insufficiently characterized. The use of hiPSC-derived cardiomyocytes provides a valuable opportunity to study the function of TOP2B in a controlled human cellular context. We have shown that iPSC deleted for TOP2B can be differentiated into cardiomyocytes, as previously reported [14]. We have extended this by carrying out a transcriptomics analysis to determine the differential gene expression in the undifferentiated and differentiated cells, both with and without TOP2B and this data has been deposited in gene expression omnibus to enable other researchers to access to this data. Four biological replicates were sequenced where RNA was extracted from independently differentiated wells of BKO iPSC clone 18D. However, the four replicas were derived from the same clonal line, which represents a potential limitation of this study. Although the BKO iPSC could differentiate efficiently and reproducibly into beating cardiomyocytes this took slightly longer than the WT iPSC suggesting some differences in the differentiation of WT and BKO into CM. Consistent with the observed CM phenotype (sheets of beating cells), both WT CM and BKO CM exhibited induced cardiomyocyte marker genes and silencing of pluripotency marker genes. However, there were significant gene expression changes between the WT CM and BKO CM. Some of the genes affected are probably direct targets of TOP2B whilst others may reflect gene expression changes due to the altered regulation of genes targeted by TOP2B.

Importantly, we showed that gene expression changes after a short exposure of WT CM cells to topobexin, a new selective TOP2B catalytic inhibitor partially phenocopies the BKO genotype. The availability of a small molecule inhibitor with selectivity for TOP2B opens the possibility of using WT CM plus or minus topobexin to determine the mechanism by which reduction of TOP2B activity mediates cardio protection following anthracycline exposure. Topobexin has been shown to protect against cardiac damage caused by anthracyclines [16] paving the way for development of TOP2B specific drugs for cardio protection. In addition, the availability of a small molecule selective TOP2B catalytic inhibitor that phenocopies a knockout will facilitate studies of the role of TOP2B in many different cell lines without the need to produce a gene knockout.

Together, our findings demonstrate that hiPSC-derived cardiomyocytes lacking TOP2B, or treated with a selective catalytic inhibitor, exhibit characteristic cardiac gene expression, while also revealing distinct transcriptional alterations associated with TOP2B loss or inhibition. This system enables the study of TOP2B function in a human cardiac context and offers a flexible platform for investigating how its modulation impacts cardiomyocyte biology. This may be used in future studies on cardiac development, epigenetic regulation, and cardiotoxicity mechanisms, including those induced by chemotherapeutic agents. However, such studies were beyond the scope of this report.

## Data availability

RNA seq data is available through GEO (https://www.ncbi.nlm.nih.gov/gds), accession GSE262148.

## CRediT authorship contribution statement

Conceptualization: VK, AJ, MK, CA; Methodology: VK, AJ, MK, JK, AC, CY, IC; Validation: IC, JC; Formal analysis: IG,JC; Investigation; VR, IC, AJ, MK; Resources: GK,IM,JR,AC,CY,ML,LA, Data Curation: IG,JC; Writing - Original Draft: VK,IC,AJ,TS,CA,MS; Writing - Review & Editing: VK,IC,AJ,TS,CA,JR,MS; Visualization: VK,IG,CA,, Supervision: CA,IC,AJ,TS,, Project administration: VK, AJ, IC, CA, Funding acquisition: TS, JR, MS, CA.

## Acknowledgements

We thank Nicola Curtin and Hannah Smith for assistance and support.

## Ethics Statement

Human iPSCs were derived from unaffected fibroblasts purchased from Lonza a commercial supplier. Ethical permission for the use of the iPSC line was not required by our institution since Lonza have an established ethical governance framework.

