## Supplementary Figure 1-3 for "Analysis of Gene Expression Changes upon Topobexin Treatment and TOP2B-knockout in hiPSC derived cardiomyocytes"

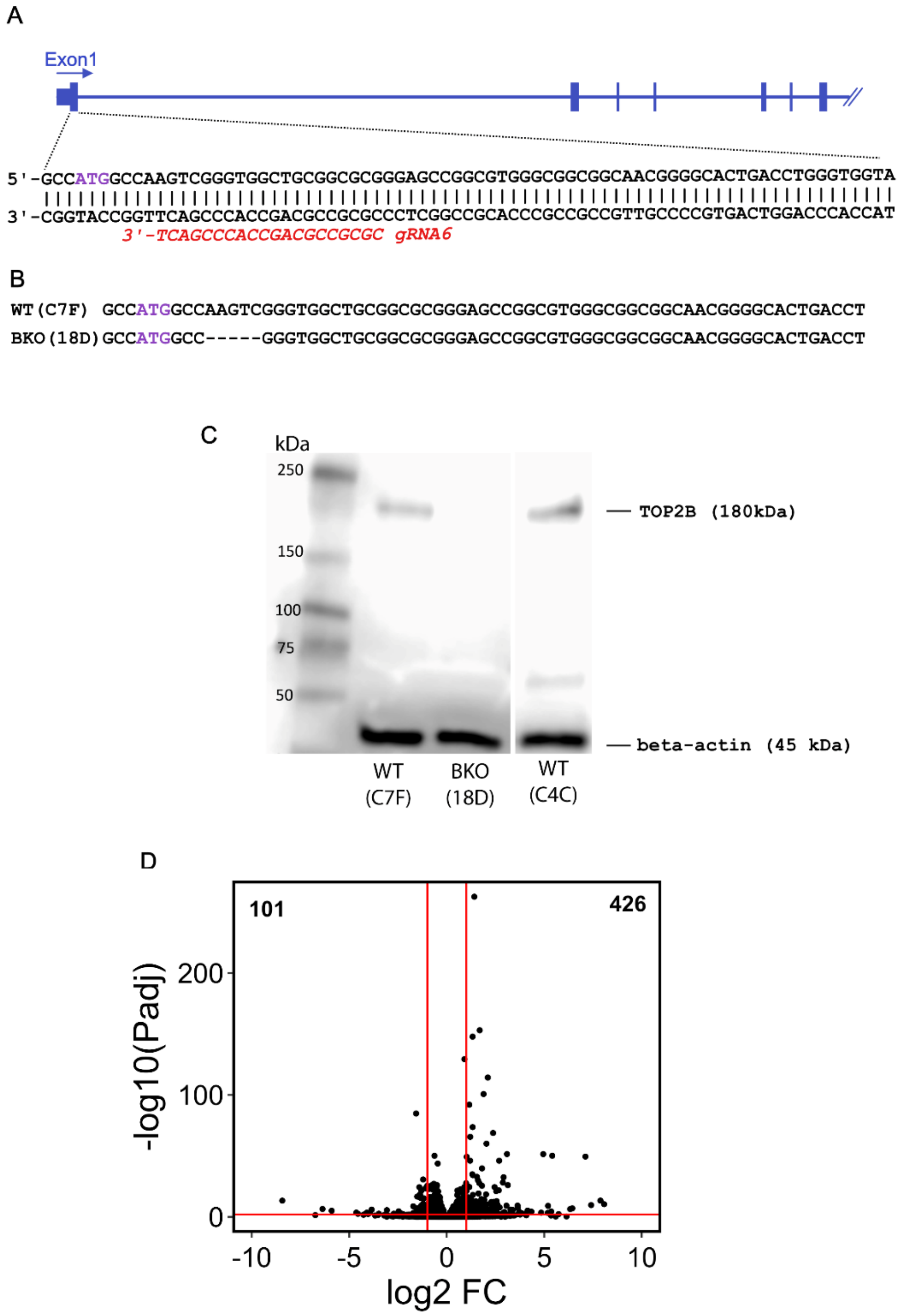


**Figure S1. CRISPR/Cas9 silencing of TOP2B in human iPSC cells. (A)** (Upper) 5'-region of the TOP2 locus illustrating the first seven exons. (Lower) DNA sequence on the 5'-coding region of exon 1 with silencing gRNA sequence shown in red. (B) DNA sequences of WT (clone CF7) and mutated (TOP2B null, BKO) clone 18D used in this study. **(C)** Western blot confirming the absence of TOP2B expression in clone 18D. **(D)** Volcano plot illustrating the transcriptional changes associated with TOP2B gene inactivation in iPSC cells. Genes upregulated in 18D (BKO, TOP2B null) cells are to the right and downregulated genes to the left. The cut-offs (red lines) are *Padj* < 0.05, $Log_2$ Fold change in gene expression >1 or <-1.

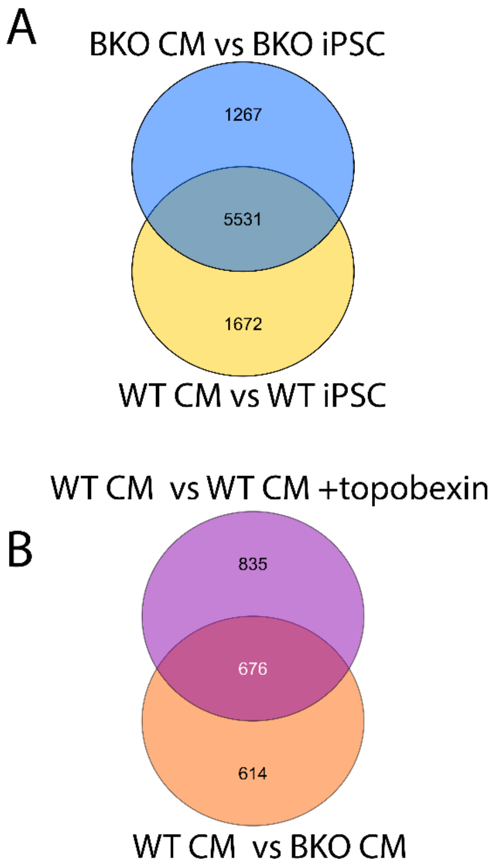


**Figure S2. (A)** Venn diagram comparing genes that change in expression in WT and TOP2B null cells (BKO) during differentiation to CM. **(B)** Venn diagram comparing differences in gene expression in WT vs TOP2B null CM cells and in WT CM vs WT CM treated with topobexin. Cut-offs are gene expression change of 2 fold or greater (in either direction) and *Padj* <0.05.

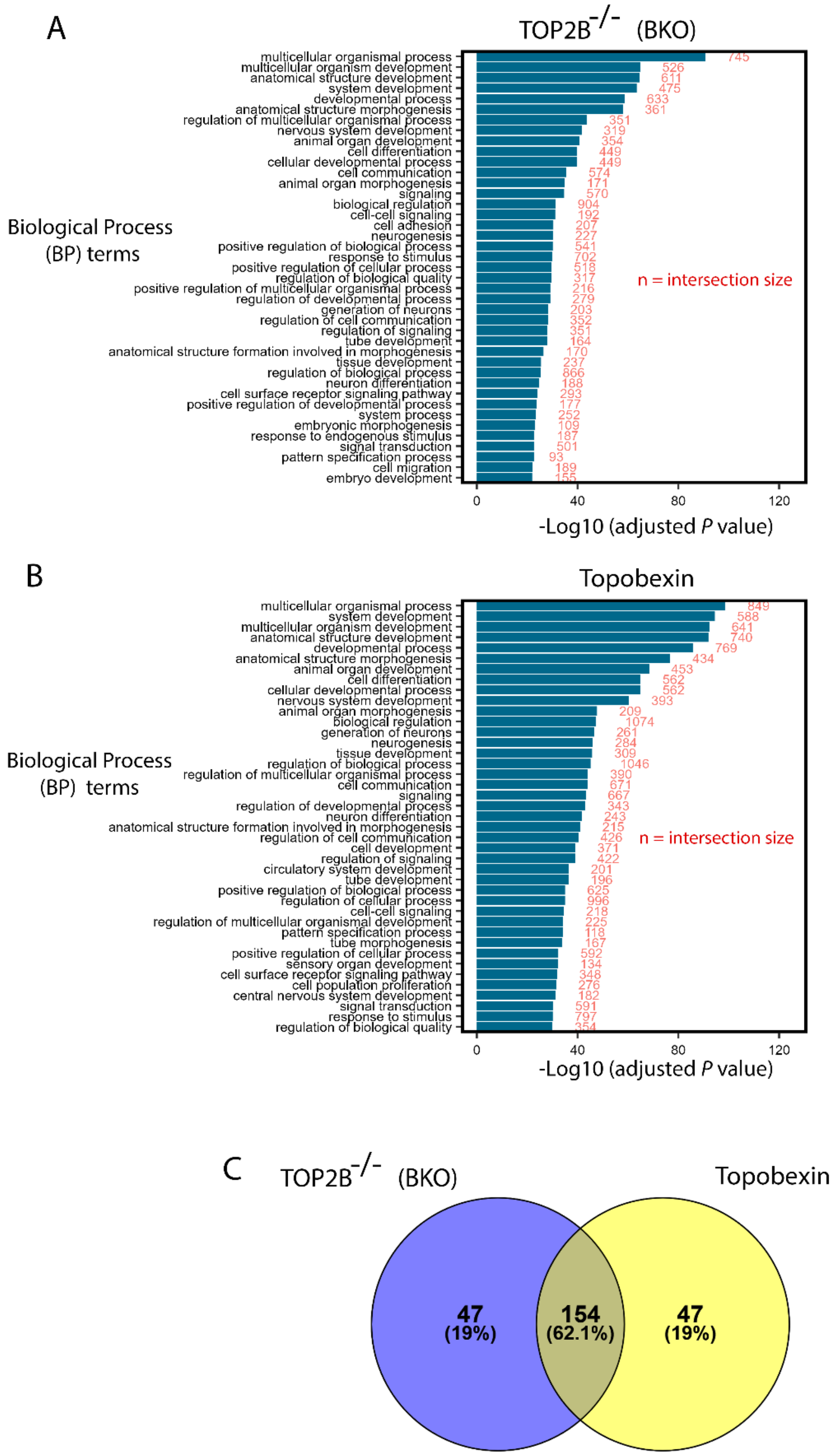


**Figure S3. Gene set enrichment analysis.** Sets of transcripts whose abundance changed (>=2 fold) in either TOP2B$^{-/-}$ (BKO) CM or topobexin-treated CM compared to untreated WT CM were examined by g:GOSt functional enrichment analysis (https://biit.cs.ut.ee/gprofiler/gost). The top 41 Biological Pathway (BP) terms for **(A)** TOP2B$^{-/-}$ (BKO) CM and **(B)** topobexin-treated CM were plotted by negative $\log_{10}$(adjusted P value). Numbers in red correspond to the intersection size. **(C)** Venn diagram illustrating the degree of overlap between the top 200 (by statistical significance) BP terms highlighted for each condition.
